# p38δ genetic ablation protects female mice from anthracycline cardiotoxicity

**DOI:** 10.1101/2020.03.02.973347

**Authors:** Sharon A George, Alexi Kiss, Sofian N Obaid, Aileen Venegas, Trisha Talapatra, Chapman Wei, Tatiana Efimova, Igor R Efimov

**Affiliations:** Department of Biomedical Engineering, The George Washington University, Washington, DC, USA; Department of Anatomy and Cell Biology, The George Washington University School of Medicine and Health Sciences, Washington, DC, USA; The George Washington Cancer Center, Washington, DC, USA; Department of Dermatology, The George Washington University School of Medicine and Health Sciences, Washington, DC, USA

**Keywords:** Doxorubicin, p38 MAPK, p38δ/MAPK13, sex differences, cardioprotection

## Abstract

**BACKGROUND:** The efficacy of an anthracycline antibiotic doxorubicin (DOX) as a chemotherapeutic agent is limited by dose-dependent cardiotoxicity. DOX is associated with activation of intracellular stress signaling pathways including p38 MAPKs. While previous studies have implicated p38 MAPK signaling in DOX-induced cardiac injury, the roles of the individual p38 isoforms, specifically, of the alternative isoforms p38γ and p38δ, remain uncharacterized.

**OBJECTIVES:** To determine the potential cardioprotective effects of p38γ and p38δ genetic deletion in mice subjected to acute DOX treatment.

**METHODS:** Male and female wild-type (WT), p38γ^-/-^, p38δ^-/-^ and p38γ^-/-^δ^-/-^ mice were injected with 30 mg/kg DOX and their survival was tracked for ten days. During this period cardiac function was assessed by echocardiography and electrocardiography and fibrosis by PicroSirius Red staining. Immunoblotting was performed to assess the expression of signaling proteins and markers linked to autophagy.

**RESULTS:** Significantly improved survival was observed in p38δ^-/-^ female mice post-DOX relative to WT females, but not in p38γ^-/-^ or p38γ^-/-^δ^-/-^ male or female mice. The improved survival in DOX-treated p38δ^-/-^ females was associated with decreased fibrosis, increased cardiac output and LV diameter relative to DOX-treated WT females, and similar to saline-treated controls. Structural and echocardiographic parameters were either unchanged or worsened in all other groups. Increased autophagy, as evidenced by increased LC3-II level, and decreased mTOR activation was also observed in DOX-treated p38δ^-/-^ females.

**CONCLUSIONS:** p38δ plays a crucial role in promoting DOX-induced cardiotoxicity in female mice by inhibiting autophagy. Therefore, p38δ targeting could be a potential cardioprotective strategy in anthracycline chemotherapy.

**NEW AND NOTEWORTHY:** This study for the first time identifies the roles of the alternative p38γ and p38δ MAPK isoforms in promoting DOX-cardiotoxicity in a sex-specific manner. While p38γ systemic deletion did not affect DOX-cardiotoxicity, p38δ systemic deletion was cardioprotective in female but not in male mice. Cardiac structure and function were preserved in DOX-treated p38δ-/- females and autophagy was increased.

## INTRODUCTION

The anthracycline antibiotic doxorubicin (DOX) is a highly potent chemotherapeutic agent that is widely used for the treatment of many different types of cancers such as sarcoma, lymphoma, leukemia, and breast cancer. However, the efficient use of DOX is limited by the development of dose-dependent, life-threatening cardiomyopathy leading to congestive heart failure, which can occur within a year (early-onset), or several years or even decades (late-onset) after DOX treatment (5, 28, 39). This poses a unique problem because not only are heart disease and cancer listed as the two leading causes of mortality in the United States (National Center for Health Statistics), but treatment of one condition contributes to the prevalence of the other.

The pathophysiological mechanisms of DOX-induced cardiotoxicity remain incompletely understood. Accumulating evidence implicates genotoxic stress associated with DOX-mediated induction of double-strand DNA breaks through inhibition of topoisomerase 2β (48, 50), oxidative stress due to increased generation of reactive oxygen species (ROS) and antioxidant depletion (12, 13), inflammation (11, 45), as well as mitochondrial and autophagy dysregulation (8, 15, 21), in DOX-induced cardiac injury, leading to cardiomyocyte dysfunction and cell death.

DOX-induced systemic inflammation and ROS generation activate stress signaling pathways in multiple cell types, including cardiomyocytes and immune cells, by upregulating and activating toll-like receptors (TLRs). TLRs are involved in cardiac stress response and inflammatory signaling, and substantially contribute to the pathology of DOX-induced cardiac injury (23, 31, 33). In cardiac myocytes, activation of the TLR signaling leads to downstream activation of the p38 mitogen activated protein kinase (MAPK) family that controls adaptive responses to stresses in mammalian cells (23).

p38 MAPK isoforms p38α, p38β, p38γ, and p38δ have tissue-specific expression patterns and context-dependent functions (6). The redundant, distinct and even opposing functions of each p38 isoform have been reported (25, 49). p38α and p38β, referred to as the conventional p38 isoforms, have been studied more extensively, while the specialized functions of p38γ and p38δ, known as the alternative p38 isoforms, are less understood, in part, due to the lack of inhibitors specific to these isoforms (7).

All four p38 MAPK isoforms are expressed in the heart (6, 25, 49). p38α and p38β appear to play opposite roles in cardiac apoptosis and cardiac hypertrophy regulation (46), while p38γ and p38δ have been reported to control cardiac growth by promoting mammalian target of rapamycin (mTOR) activation through phosphorylation and degradation of the mTOR inhibitor protein DEPTOR (9).

Although p38 MAPK signaling has been implicated in DOX-induced cardiotoxicity, the roles of the individual p38 isoforms in regulating the mechanisms underlying the cardiotoxic side effects of DOX remain thus far uncharacterized. Thus, previous approaches, such as a simultaneous co-inhibition of either a subset (for example, using a pharmacologic inhibitor specific for p38α/p38β (3)), or all four p38 isoforms (for instance, using a dominant-negative mutant approach (41)) confounded the analysis of the reported phenotypes. Moreover, the roles of p38γ and p38δ in anthracycline cardiotoxicity have not been explored previously.

In this study, we investigated the roles of the alternative p38 isoforms p38γ and p38δ in DOX-induced cardiotoxicity using systemic p38γ, p38δ or p38γ/p38δ knockout mice.

## METHODS

All animal protocols were approved by the Institutional Animal Care and Use Committee at The George Washington University and conform to the guidelines of the National Institutes of Health Guide for the Care and Usage of Laboratory Animals. Raw data files will be made available upon request.

### Mouse Strains

All mice used in this study were on a C57BL/6J background. Wild type (WT) mice were ordered from the Jackson Laboratory (Stock No: 000664). Generation of mice with systemic deletion of p38γ and p38δ was previously reported (35). Genotyping was carried out as described (35). Mice with systemic deletion of p38γ and p38δ were bred in-house to generate mice that lacked p38γ (p38γ^-/-^), p38δ MAPK (p38δ^-/-^) and both p38γ and p38δ (p38γ^-/-^δ^-/-^). The efficient knockdown of p38γ and p38δ was confirmed by western blotting (Supplemental Figure 5B). Male and female mice were used in experiments at ∼15 weeks of age. An overview of the mouse genotypes used in this study and the treatments they were subjected to is illustrated in Figure 1A.

**Figure 1:**
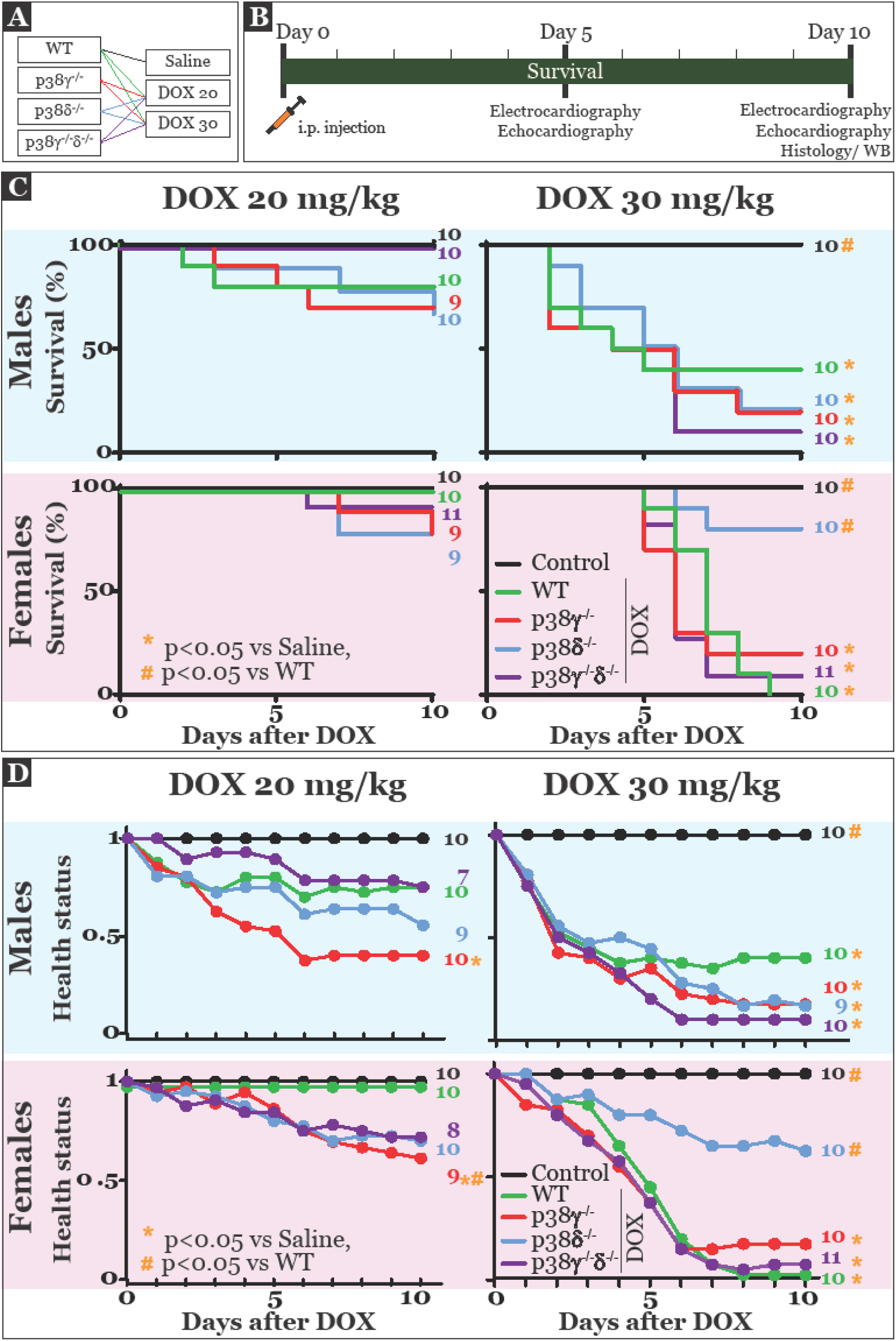
Increased survival rate and preserved health status in female p38δ^-/-^ mice relative to female WT mice ten days after DOX treatment. **A)** The genotypes of mice used in this study and the treatments they were subjected to are detailed. **B)** Timeline of the experiments. Functional data from the surviving mice were acquired on Days 5 and 10 post-DOX injection, and hearts were collected for histological and protein expression assessment on Day 10. **C)** Survival rates in male and female mice (upper and lower panels, respectively) treated with 20 or 30mg/kg DOX (left and right panels, respectively). **D)** Health status was tracked in male and female mice (upper and lower panels, respectively) treated with 20 or 30mg/kg DOX (left and right panels, respectively) over the 10 day survival period using an arbitrary health scoring system as detailed in the Methods section.

### Survival Protocol

The study protocol is illustrated in Figure 1B. Mice were injected intraperitoneally (i.p.) with either 20 mg/kg or 30 mg/kg of DOX, or saline (vehicle control), and the survival of the mice and their health status were tracked over the next 10 days. Each day, mice were assigned a number between 1 and 0 to indicate their health status. A health score of 1 indicated healthy mice with no visible signs of distress, 0.75 indicated slight impairment such as slight hunch in posture, 0.5 indicated sick mice that were hunched and showed slow movement, 0.25 indicated moribund mice and 0 indicated dead mice. During, the 10-day survival period cardiac function was assessed on Days 5 and 10 by echocardiography and electrocardiography as indicated in Figure 1B. On Day 10, surviving mice were euthanized and their hearts were collected following thoracotomy and either fixed in 4% paraformaldehyde for histological assessment or flash frozen in liquid nitrogen for protein extract isolation for western blotting.

### Electrocardiography (ECG)

On Days 5 and 10 post-DOX injection, ECG was recorded from surviving mice in conscious state using the non-invasive emka ECG tunnel recording device (emka TECHNOLOGIES, Paris, France). Briefly, mice were guided into a mouse size-specific tunnel over a platform fitted with four ECG electrodes. The mouse was positioned on the platform such that the four paws were in contact with the electrodes and ECG was recorded using the IOX software. The recorded signals were then analyzed using a custom Matlab program to determine P wave duration (P), P-R interval (PR), QRS duration (QRS), QT interval corrected for HR (QTc) and R-R interval (RR).

### Echocardiography

After ECG measurements, mice were anesthetized by inhalation of 2.5% isoflurane at 1 ml/min oxygen flow using EZ anesthesia machine. Once unresponsive, the mice were transferred to an imaging stage fitted with ECG electrodes to measure heart rate (HR). Isoflurane level was adjusted between 1 and 2.5% to maintain a HR of ∼400 bpm, except in sick mice where HR was reduced irrespective of anesthesia. Chest hair was removed and the ultrasound gel (Aquasonics, Clear) was applied on the chest, left of the sternum. M-mode echocardiography of the left ventricle (LV) was performed using Vevo 3100 system (VisualSonics/Fujifilm) and the acquired data were analyzed using VevoLAB2.1.0. ejection fraction (EF), cardiac output (CO), stroke volume (SV), fractional shortening (FS), LV posterior wall thickness during systole (LVPWs) and diastole (LVPWd) and LV end systolic and diastolic diameters (LVESD and LVESD, respectively) were measured.

### Histology

At the end of the survival period, mice were euthanized and hearts were collected and fixed in 4% paraformaldehyde for 48 hours and then transferred to 70% ethanol. Hearts were then paraffin embedded, sectioned onto slides in the cross-sectional orientation and stained using the PicroSirius Red as per manufacturer instructions (Abcam), to assess collagen deposition. Slides were then imaged using a light microscope and three images each from the LV and right ventricle (RV), from the base, mid and apex regions, were acquired. The images were analyzed using a custom Matlab program to determine the percentage of area occupied by collagen (red) in both the LV and RV.

### Western Blotting

Hearts isolated from mice were rinsed in PBS to remove blood and flash frozen in liquid nitrogen. Ventricular tissue was cut from these frozen sections and lysed in RIPA lysis buffer (Sigma, 89901) supplemented with protease and phosphatase inhibitor tablets (Thermofisher, A32955). Total protein concentration was determined using a BCA kit (Thermo Scientific, 23227) following manufacturer instructions. Electrophoresis was performed to separate the proteins on a 4-15% Biorad Criterion gel (Biorad, 5671084) and transferred to PVDF membrane. Membranes were then blocked using Odyssey Blocking buffer (Licor, 927-5000) and incubated overnight in primary antibodies at 4°C. Membranes were then washed and incubated with secondary antibodies for 2 hours at room temperature. Membranes were washed again and imaged using a Licor FC imager and protein expression was measured using Image Studio Lite software. All protein expressions are normalized to GAPDH to account for loading errors. Primary antibodies used in this study include p38α (Sigma-Aldrich, M0800), p38γ (Cell Signaling, 2307), p38δ (MRC PPU, University of Dundee), mTOR (Cell Signaling, 2972), phospho-mTOR Ser2488 (Cell Signaling, 5535), Akt (Cell Signaling, 4691), phospho-Akt Th308 (Cell Signaling, 13038), DEPTOR (Novus Biologicals, NBP1-4967), LC3 (Origene, TA301542) and GAPDH (Abcam, ab9484). Secondary antibodies used in this study include IRDye 680 Donkey anti-mouse (Li-cor, 926-68022) and IRDye 800 Donkey anti-rabbit (Licor, 926-32213).

### Statistics

All data are reported as mean ± standard deviation unless stated otherwise. Sample sizes are listed in the figures. Log rank tests were performed to determine statistically significant differences in the mouse survival data. For all other data sets, significance was determined by 1-way ANOVA with genotype as the single factor and Student’s t-tests were performed for post-hoc analysis. A p value less than 0.05 was considered statistically significant.

## RESULTS

### Female p38δ^-/-^ mice are protected against DOX-induced mortality and morbidity

Activation of p38 MAPK in hearts of WT male mice acutely treated with DOX has been previously demonstrated (3). We first examined whether p38 MAPK signaling pathway was activated in WT female mice treated with DOX at 30 mg/kg of body weight by i.p. injection. As shown in Supplemental Figure 1, we detected a significant increase in phosphorylation/activation of p38 MAPK as well as of the upstream activators (MKK3 and MKK6) and downstream effectors (p53 and HSP27) of the p38 MAPKs in DOX-treated versus saline-treated WT mouse hearts. This data indicates a sustained (lasting for at least 5 days) activation of p38 MAPK signaling pathway by DOX treatment in our mouse model. Of note, all four p38 isoforms could potentially contribute to this activation.

Next, the roles of p38γ and p38δ, the two relatively understudied p38 isoforms that are abundant in heart (9) (Supplemental Figure 5B), in DOX-induced cardiotoxicity were investigated. WT, p38γ^-/-^, p38δ^-/-^, and p38γ^-/-^δ^-/-^ male and female mice were treated with DOX at 20 or 30 mg/kg of body weight (DOX20 and DOX30, respectively) by i.p. injection, as outlined in Figure 1A-B. The survival and health of mice were acutely tracked over the next 10 days. While all mice treated with saline (control, black line, Figure 1C) survived the acute survival period, those treated with DOX had varying responses depending on their genotype and sex, as well as the dose of DOX.

The survival rates were not significantly affected by treatment with DOX20 relative to control in any of the experimental groups (Figure 1C, left panels). In contrast, the acute survival was significantly reduced in all groups treated with DOX30 relative to control, except in female p38δ^-/-^ mice (Figure 1C, right panels). Among WT mice treated with DOX30, only 40% of males and no females survived the 10 day period post-DOX. In the p38γ^-/-^ mice, 20% survival rate and in the p38γ^-/-^δ^-/-^ mice, 10 % survival rate was observed, in both males and females. Remarkably, while only 20% of DOX30-treated male mice survived the acute survival period, 80% of female p38δ^-/-^ mice survived. These findings suggest that p38δ is a crucial determinant of DOX cardiotoxicity in female mice.

Notably, significant sex-dependent differences in survival were observed as early as 5 days post-DOX injection. As shown in Supplemental Table 1, the 5-day survival rates of DOX30-treated male WT, p38δ^-/-^, and p38γ^-/-^δ^-/-^ mice were significantly lower compared with their respective female counterparts.

To determine if DOX-induced morbidity correlated with mortality, we additionally tracked the health status of the DOX-treated mice over the 10-day survival period, as detailed in the Methods section (Figure 1D). Comparing the survival data and the health status (Figures 1C vs 1D), demonstrates that although the survival rates were not significantly reduced in mice treated with DOX20, the surviving mice were of poor health, specifically male and female p38γ^-/-^ mice (Figure 1D, left panels). Among mice treated with DOX30, survival rates correlated with health status across genotypes. Importantly, the higher survival rate seen in the female p38δ^-/-^ group correlated with better health status similar to controls (Figure 1D, lower right panel).

### DOX-treated female p38δ^-/-^ mice exhibit preserved cardiac function

Cardiac function deteriorates due to DOX-induced cardiotoxicity. We next assessed cardiac mechanical function in DOX30-treated WT, p38γ^-/-^, p38δ^-/-^, and p38γ^-/-^δ^-/-^ male and female mice by M-mode echocardiography. Data obtained 5 days post-DOX treatment are summarized in Figure 2 and Table 1. Day 10 assessment summary for the surviving mice is included in the Supplemental Figure 2.

**Table 1.**
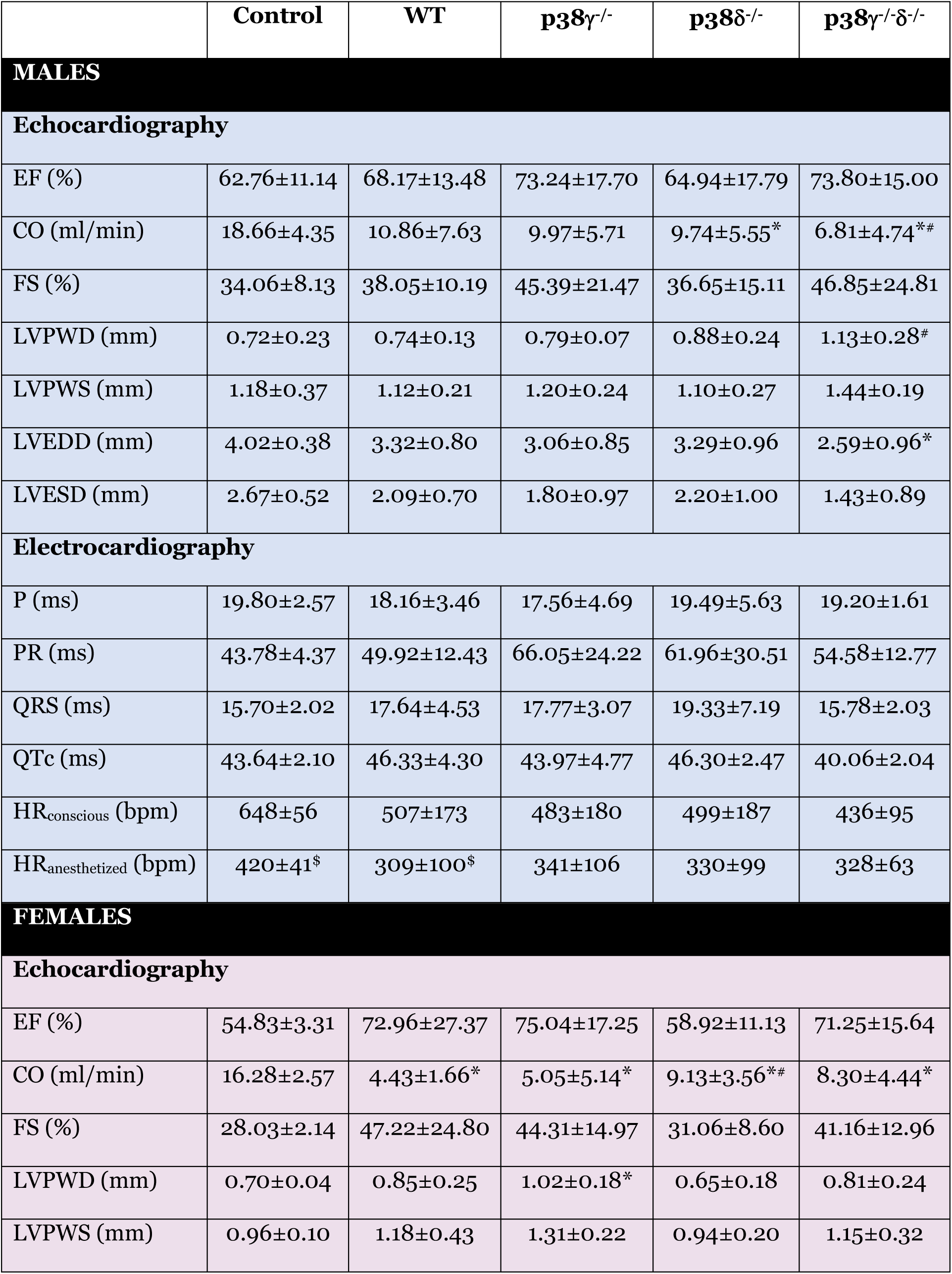
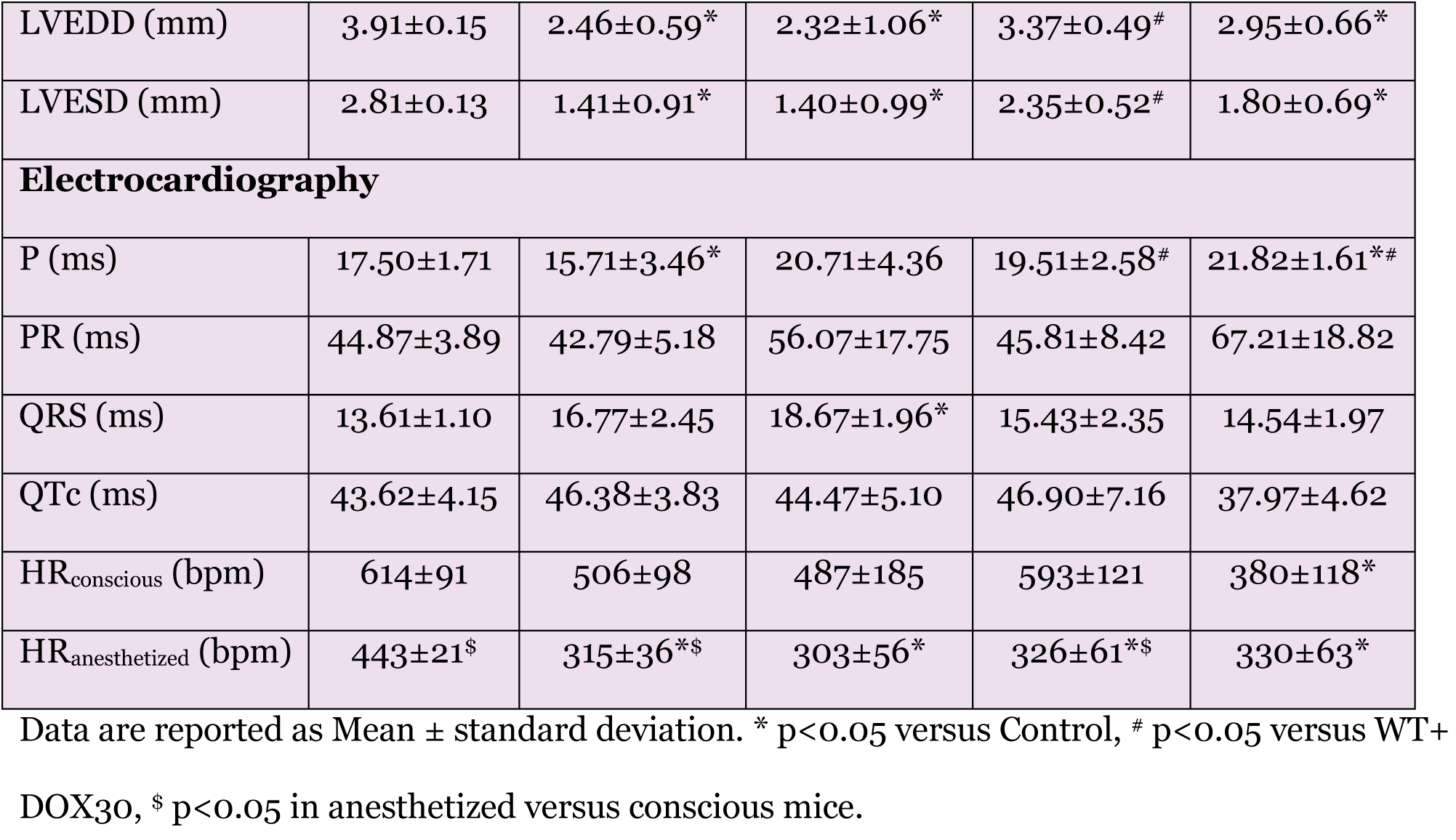
Summary of Cardiac Functional Data.

**Figure 2.**
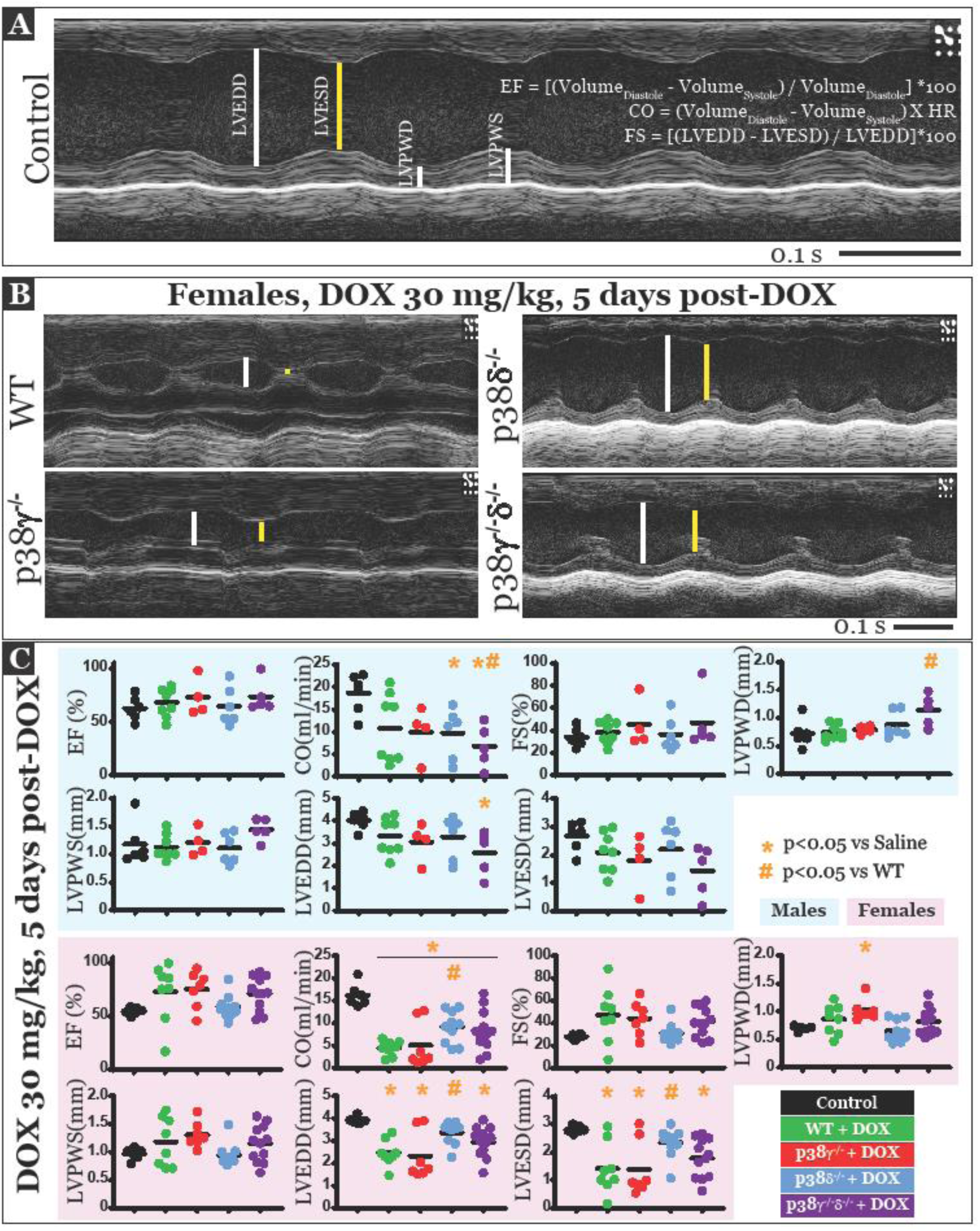
Cardiac mechanical function and LV structure are preserved in DOX-treated female p38δ^-/-^ mice. **A)** Representative echocardiogram from a control (saline-treated) female mouse with the description of the specific parameters measured. **B)** Representative echocardiograms from female WT, p38γ^-/-^, p38δ^-/-^ and p38γ^-/-^δ^-/-^ mice 5 days after DOX injection. **C)** The indicated parameters of cardiac mechanical function were measured in male (upper blue panel) and female (lower pink panel) mice treated with 30mg/kg DOX, 5 days post-treatment. EF: Ejection Fraction, CO: Cardiac Output, FS: Fractional Shortening, LVPWD, LVPWS: Left Ventricular Posterior Wall Thickness during diastole/systole, LVEDD, LVESD: Left Ventricular End Diastolic/Systolic Diameter.

Ejection fraction (EF) and fractional shortening (FS) were not significantly altered in the DOX30-treated male or female mice of any of the genotypes examined. However, a major indicator of DOX-cardiotoxicity in our mouse model was a significant reduction in cardiac output (CO). All female mice treated with DOX had significantly lower CO relative to control mice (Figure 2C). Although CO was reduced in female p38δ^-/-^ mice (improved survival rate) relative to saline-treated controls, CO in this group was also significantly enhanced relative to DOX30-treated WT mice, indicating a protection of cardiac function in the former group. In male mice, CO was reduced in all DOX30-treated mice relative to controls, however, this reduction was only significant in p38δ^-/-^ and p38γ^-/-^p38δ^-/-^ mice. CO in DOX30-treated male WT and p38γ^-/-^ mice was not significantly different from controls (p=0.09 and 0.05, respectively) possibly due to larger variations in CO response to DOX treatment in these mice. Additionally, DOX-treated male p38γ^-/-^p38δ^-/-^ mice had significantly reduced CO compared to DOX30-treated male WT mice (Figure 2C). High mortality rates in male DOX30-treated mice precluded the assemblage of a larger sample size in these groups 5 days post-DOX treatment. To address this limitation, additional experiments were performed at an earlier time point (Day 3 post-DOX) in male WT mice treated with DOX30. As shown in Supplemental Figure 3, a significant reduction in CO was observed in DOX30-treated male WT mice relative male control mice, while no changes in EF or FS were detected.

Next LV parameters such as wall thickness and diameter were measured to assess heart failure due to DOX treatment. LV wall thickness (LVPWD) was increased or hypertrophic in DOX30-treated female p38γ^-/-^ mice, while LV diameter was significantly reduced in DOX30-treated female WT, p38γ^-/-^ and p38γ^-/-^δ^-/-^ mice at the end of systole and diastole (LVEDD and LVESD). Importantly, these detrimental effects were not observed in DOX30-treated female p38δ^-/-^ mice which exhibited improved survival. Among male mice, LVPWD was increased and LVEDD was decreased in p38γ^-/-^δ^-/-^ mice relative to DOX30-treated WT and saline-treated controls, respectively (Figure 2C).

To summarize the echocardiographic analysis, cardiac mechanical function and LV wall structure were significantly perturbed by DOX treatment in male and female mice (Figure 2 and Supplemental Figure 3). However, these parameters were preserved in DOX30-treated female p38δ^-/-^ mice, in line with the improved health and survival rate observed in this group.

### ECG parameters are preserved in DOX-treated female p38δ^-/-^ mice

The effects of DOX treatment on cardiac electrophysiology were determined by recording ECGs in conscious mice. Illustrated and summarized in Figure 3 and Table 1 are ECG traces and parameters from mice treated with DOX30, 5 days post-DOX. Day 10 ECG data are provided in Supplemental Figure 4.

**Figure 3.**
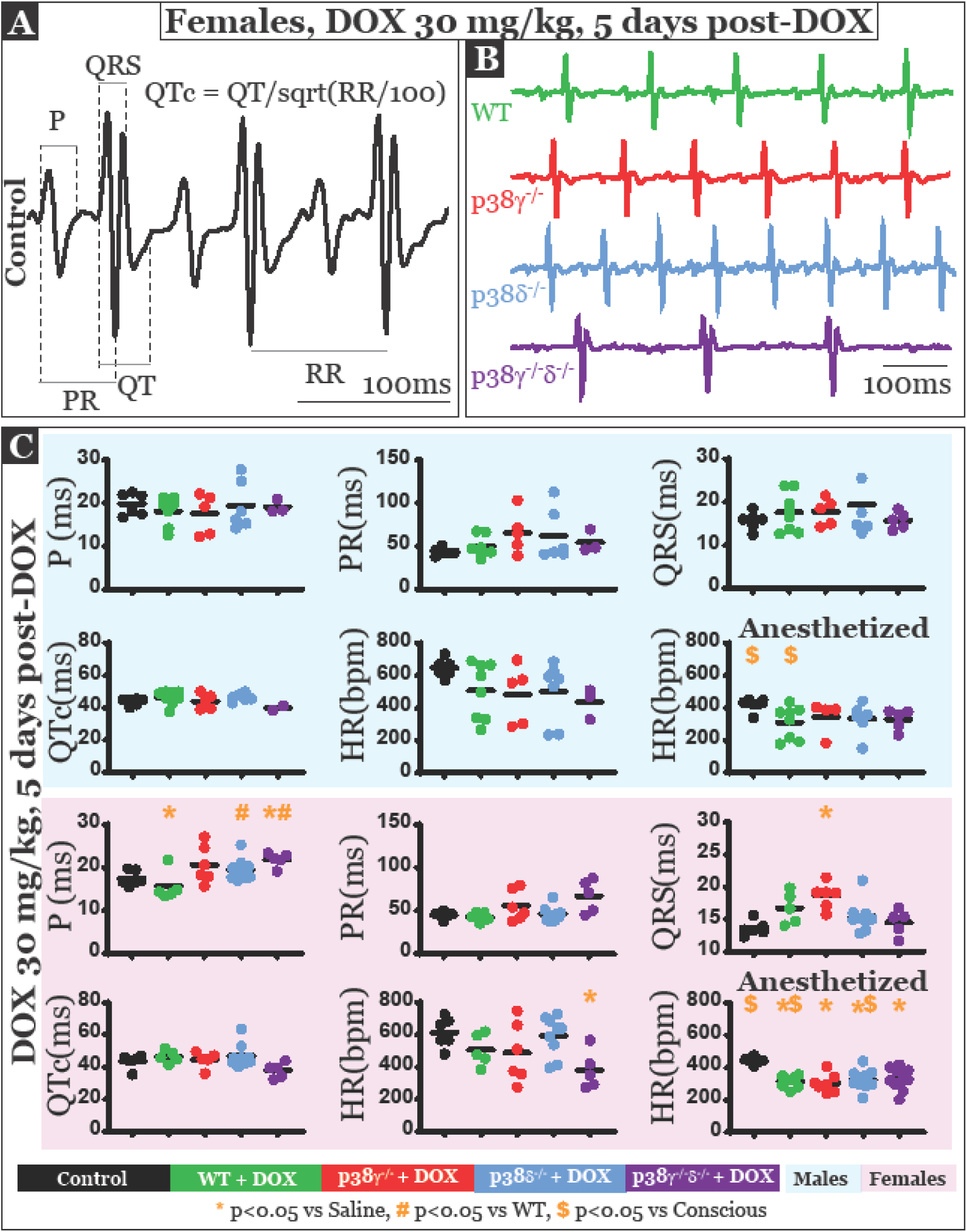
Electrocardiogram (ECG) parameters are preserved in DOX-treated female p38δ^-/-^ mice. **A)** Representative ECG from a female control mouse with the description of the specific parameters measured. **B)** Representative ECGs from female WT, p38γ^-/-^, p38δ^-/-^ and p38γ^-/-^δ^-/-^ mice. **C)** ECG parameters such as the P wave interval (P), P-R interval (PR), QRS duration (QRS), Q-T interval corrected for heart rate (QTc) and heart rate (HR) were measured in conscious male (upper blue panel) and female (lower pink panel) mice treated with 30 mg/kg DOX 5 days post-treatment. HR in anesthetized mice was also measured for comparison.

Once again, modulation of cardiac function by DOX was measurable only in female mice. Specifically, P interval was prolonged in WT and p38γ^-/-^δ^-/-^ mice treated with DOX30 versus control, and QRS duration prolongation was observed in DOX30-treated p38γ^-/-^ mice versus control. These data indicate impaired or slow conduction of electrical impulses in the atria and ventricles, respectively. Additionally, HR was reduced in female p38γ^-/-^δ^-/-^ mice relative to DOX-30-treated female WT mice. Importantly, ECG parameters in female p38δ^-/-^ mice were similar to controls. No significant changes were detected in ECGs of male mice (Figure 3C and Table 1).

Lastly, HR from conscious mice was compared to anesthetized mice. While DOX30-treatment did not modulate HR in any anesthetized male mice, HR was significantly reduced in all DOX30-treated female anesthetized mice relative to control (Figure 3C and Table 1). Differential HR response to DOX treatment in conscious versus anesthetized state requires further investigation. Furthermore, significant reduction in HR between conscious and anesthetized state was detected in male and female control and WT mice as well as in female p38δ^-/-^ mice. This is consistent with previously published data that reported slowing down of HR with isoflurane anesthesia (47). The lack of significant changes in the p38γ^-/-^ and p38γ^-/-^δ^-/-^ groups could be due to a trend towards lower HR in the conscious state seen in these groups to begin with.

### Fibrosis is reduced in hearts of DOX-treated female p38δ^-/-^ and p38γ^-/-^δ^-/-^ mice

Extracellular edema and fibrosis are commonly observed during several cardiac diseases including heart failure (29, 30, 37). DOX treatment has previously been shown to induce fibrosis in the heart (19, 21). We next assessed myocardial collagen deposition in mice treated with DOX30 on Day 10 post-treatment using PicroSirius Red staining (Figure 4). Fibrosis in left and right ventricles (LVs and RVs) was evaluated separately in female mice of all groups (Figure 4B). Male mice were not included due to high mortality rate in this group at Day 10 post-DOX.

**Figure 4.**
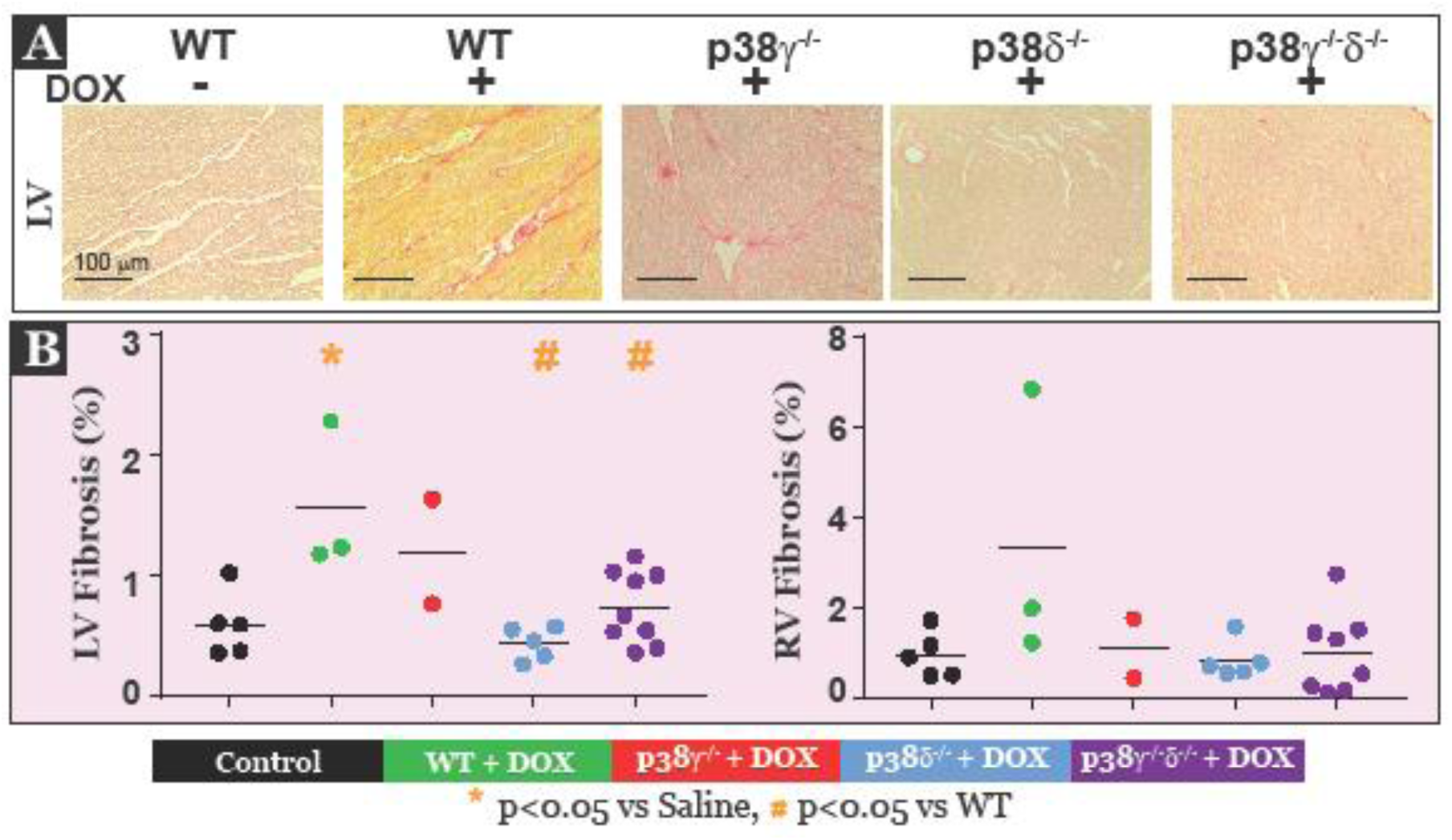
Fibrosis is reduced in left ventricles of the surviving DOX-treated female p38δ^-/-^ mice. **A)** Representative images of PicroSirius Red staining in LV of female mice of the indicated genotypes treated 30 mg/kg DOX (+) or with saline (-). Tissue samples were collected on Day 10 post-treatment. Scale bar, 100 μm. **B)** Quantification of the areas occupied by fibrosis as visualized by red positive staining (collagen) in left and right ventricles of female mice was carried out as detailed in the Methods section.

As expected, increased fibrosis, demonstrated by enhanced PicroSirius Red staining, was observed in the LV of female WT mice treated with DOX relative to controls. Notably, LV fibrosis in DOX30-treated female p38δ^-/-^ and p38γ^-/-^δ^-/-^ mice was significantly reduced relative to DOX30-treated female WT mice and was similar to control values. Interestingly, DOX treatment did not change fibrosis in the RV of mice in any of the DOX30-treated groups relative to control (Figure 4B, right panel). The observed preservation of cardiac structure in the DOX30-treated female p38δ^-/-^ mice was in keeping with the improved health and survival as well as the preservation of cardiac function in this group.

### p38δ deletion results in reduced mTOR activation and increased autophagy in hearts of DOX-treated female mice

DOX cardiotoxicity has been linked to dysregulation of autophagy in the myocardium (2, 19). mTOR signaling has been shown to regulate autophagy, specifically, to provide maladaptive feedback inhibition of DOX-induced autophagy in the heart (21). Notably, p38γ and p38δ have been implicated in the control of mTOR activity in the heart by means of phosphorylation and subsequent degradation of the mTOR inhibitory protein DEPTOR (9). We therefore examined the potential effect of the individual or combined p38δ and p38γ deletion on myocardial mTOR/Akt signaling and autophagy in DOX30-treated female mice 10 days post-DOX treatment. Male mice were not included due to high mortality rate in this group at Day 10 post-DOX.

As illustrated in Figure 5A and 5B, the expression of total mTOR protein and the phosphorylation of mTOR at Ser 2488 were significantly reduced in DOX30-treated female p38δ^-/-^ mice relative to DOX30-treated female WT mice, as assessed by immunoblotting. On the other hand, no significant changes were detected in total Akt protein expression or in Akt phosphorylation at Thr308 among the specified groups of mice included in the analysis. However, Akt phosphorylation at Ser473 was significantly reduced in female mice of all genotypes treated with DOX, relative to control (Figure 5A and 5B).

**Figure 5.**
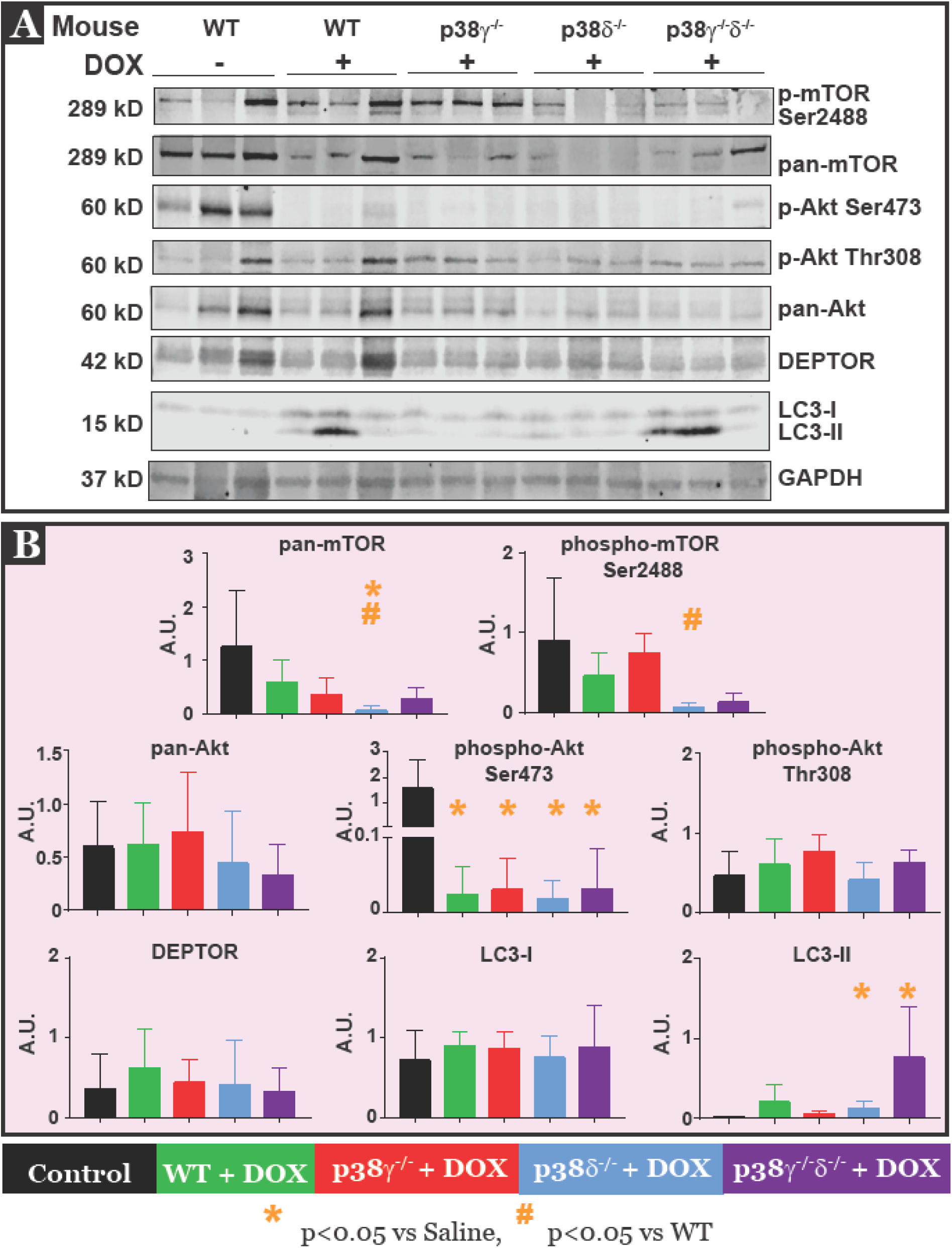
Reduced mTOR activation and increased expression of autophagy marker LC3-II in hearts of DOX30-treated female p38δ^-/-^ mice. **A)** Immunoblot and **B)** relative quantification of the specified proteins in hearts from female mice of the indicated genotypes. Lysates from the hearts of three individual mice were included per each group. The quantification data include the results of two technical repeats of the immunoblot experiment. Each protein expression was normalized to GAPDH expression in the same sample to avoid loading errors.

At variance with the previously published data (9), possibly due to different experimental settings, we did not observe an increase in the expression of DEPTOR in the hearts of female p38γ^-/-^, p38δ^-/-^, or p38γ^-/-^δ^-/-^ mice compared with their WT counterparts (Figure 5A and 5B). Finally, as shown in Figure 5B, the levels of LC3-II, a marker of autophagosome formation, were significantly increased in female p38δ^-/-^ and p38γ^-/-^δ^-/-^ mice relative to control.

These data suggest that a resistance of female p38δ^-/-^ mice to DOX cardiotoxicity could be ascribed to the loss of mTOR-dependent negative regulation of DOX-stimulated autophagic response. Interestingly, as described above, DOX30-treated female p38γ^-/-^δ^-/-^ mice exhibited protective phenotypes such as reduced fibrosis and increased autophagy marker expression even though this group had lower survival rates relative to controls 10 days post-DOX30 treatment. However, in a separate survival experiment using groups of older female mice (22 weeks of age), p38γ^-/-^δ^-/-^ females fared significantly better with DOX20 treatment, with a 100% survival rate compared with 60% survival rate in the DOX20-treated WT female cohort (Supplemental Figure 5A). An increase in the autophagy marker LC3-II was also observed in this group (Supplemental Figure 5C). This suggests that the combined deletion of both p38γ and p38δ, while beneficial at lower DOX doses, cannot sustain normal cardiac function at higher DOX doses. The pathways involved in the development of cardiotoxicity in this group will require further investigation.

## DISCUSSION

In this study we investigated the roles of stress-activated alternative p38 MAPK isoforms p38γ and p38δ in the development of cardiotoxic effects induced by treatment with the anthracycline antibiotic DOX. We report that a systemic genetic deletion of p38δ protects female mice against DOX-induced cardiotoxicity. Our findings, summarized in Figure 6, revealed that cardiac electrical and mechanical functions were preserved in female mice lacking p38δ, resulting in a significantly higher survival rate 10 days after DOX injection compared with WT female mice (80% vs 0%, p<0.05). One mechanism that could contribute to cardioprotection in this group could be increased autophagy.

**Figure 6.**
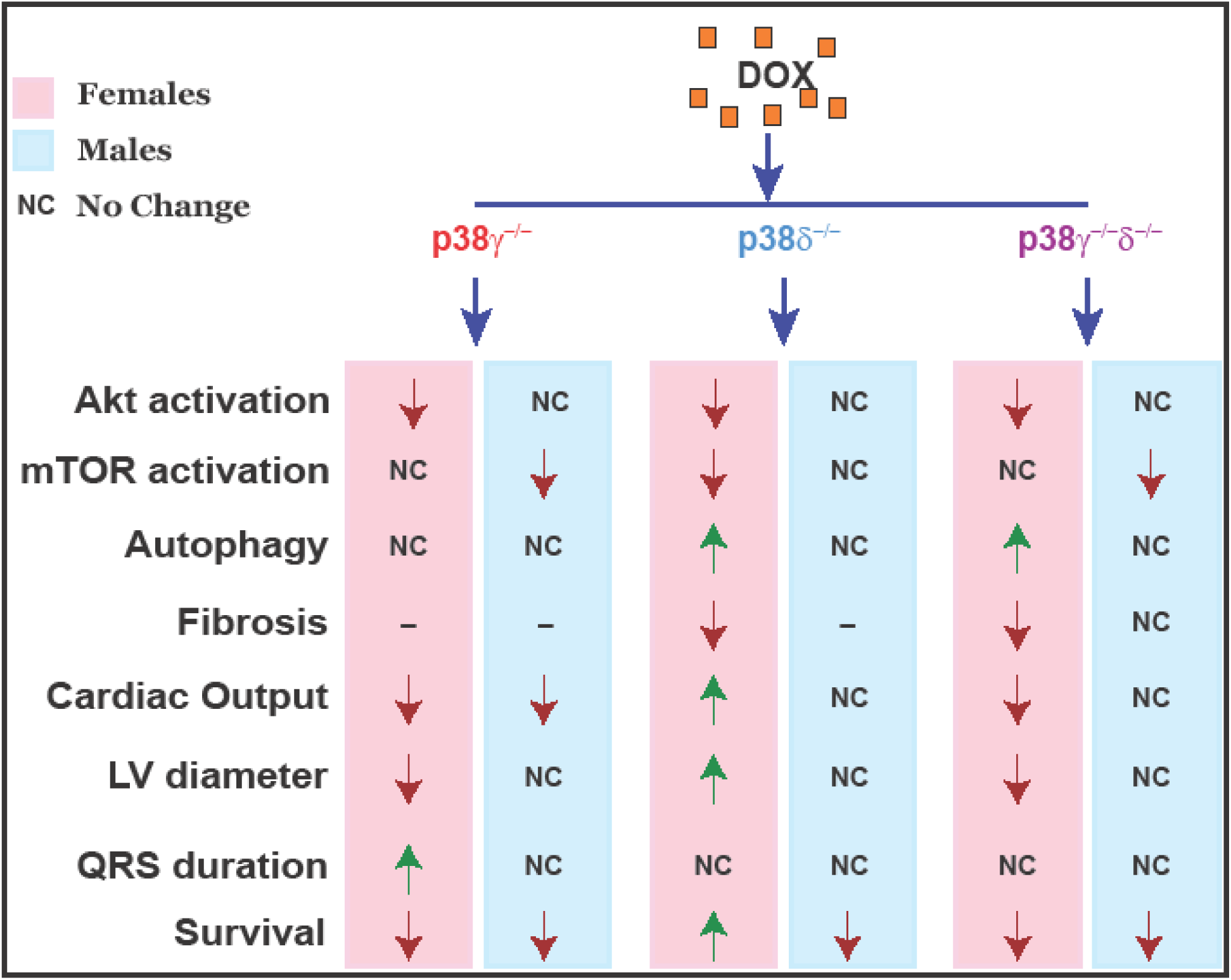
p38δ deletion increases autophagy and preserves cardiac function and survival in female mice: summary of findings.

### Sex-dependent differences in doxorubicin cardiotoxicity

One of the important novel findings of this study is the sex-dependent differences in cardiotoxicity and survival. Notably, clinical data showed that the male sex is a risk factor for adverse cardiac effects in adult patients undergoing anthracycline therapy (26). Consistently, regardless of genotype, male mice treated with DOX displayed an average of 45% survival rate 5 days post-DOX while in females groups an average of 86% mice survived this time period. The mechanisms underlying the sex-dependent differences in responses to anthracycline therapy merit further investigation.

### Preserved Ejection Fraction during DOX treatment

While it is well-established that reduced EF is a hallmark of DOX treatment, transient changes in EF especially during early phases of chemotherapy have also been reported (38, 42). No change or even improvement in EF immediately after the first cycle of chemotherapy have been reported in patients (38, 42). In our acute DOX treatment model, we observed preserved EF in all groups of mice treated with DOX at Days 3 and 5 post-DOX treatment. One possible explanation for this phenomenon could be the simultaneous modulation of both systolic and diastolic function by DOX. To elucidate this further, EF is calculated using the ratio of blood ejected from the heart during systole to the maximal LV loading during diastole. Reduced systolic ejection and diastolic loading at the same time can result in preserved EF. Of note, DOX has been reported to cause both systolic and diastolic dysfunction (4, 51).

The acute, transient changes in EF could complicate early diagnosis of DOX-induced cardiotoxicity using this metric. On the other hand, in our mouse model, CO was acutely reduced in males and female after DOX treatment suggesting that reduced CO could serve as an earlier indicator of DOX cardiotoxicity.

### Stress signaling in doxorubicin cardiotoxicity

Generation of ROS is suggested as one of the potential mechanisms underlying anthracycline-induced cardiotoxicity (12, 13). Increased ROS generation has multiple downstream effects that can contribute to development of heart failure symptoms such as mitochondrial dysfunction, altered intracellular calcium homeostasis and apoptosis among others, resulting in a significant impairment of cardiac electrical and mechanical functions. Previous *in vivo* studies have implicated p38 MAPK signaling in DOX cardiotoxicity (3, 41, 43). Most of these studies focused mainly on the conventional p38α and p38β isoforms, employing a specific inhibitor of p38α/p38β, SB203580, as a tool to delineate the contributions of these isoforms to DOX cardiotoxicity (3, 43). On the other hand, transgenic overexpression of a dominant-negative mutant form of p38α in cardiomyocytes (41) would be expected to co-inhibit all four p38 isoforms, potentially obfuscating the roles of the individual p38 isoforms given their reported diverse and at times opposing roles in controlling certain aspects of cardiac physiology and pathology (34).

In this study we focused on the contributions of the alternative p38 isoforms, p38γ and p38δ, the roles of which have not been previously examined in the context of anthracycline-induced cardiotoxicity. We report that, while p38γ deletion did not significantly modulate cardiac function or survival after DOX treatment, p38δ deletion resulted in significantly improved cardiac function and survival in DOX-treated female mice. Cardioprotection in female p38δ^-/-^ following DOX treatment was associated with decreased mTOR protein expression and activity in the hearts of mice surviving DOX treatment, and increased expression of the autophagic marker LC3-II, suggesting the potential cardioprotective role of autophagy during DOX treatment.

Importantly, in this study we implemented systemic p38γ and p38δ deletion models. This approach provides us the advantage of identifying the effects of p38γ and/or p38δ depletion/inactivation not only in the heart but also in other cell types such as immune cells which can in turn affect cardiac function. Furthermore, based on the results of this study, any potential therapy targeting p38δ would likely be administered systemically. Relevantly, our data demonstrate that systemic p38δ deletion does not affect normal physiology and survival of these mice, while providing significant cardioprotection in female mice during DOX treatment.

Finally, p38δ deletion has also been reported to result in anti-cancer effects in several mouse models of cancer development, including skin, breast, and colon cancer models (16, 17, 32, 36, 44). Thus, targeting p38δ in combination with DOX chemotherapy could have a dual benefit, potentially enhancing anti-cancer effect of DOX, while providing cardioprotection, particularly in female patients.

### Protective role of autophagy in doxorubicin cardiotoxicity

Autophagy is a housekeeping mechanism that maintains the equilibrium in the cellular environment by removing damaged or degraded components. A delicate balance in autophagy is required for efficient cellular function (10, 24). On the one hand, decrease in autophagy can result in the accumulation of degraded products within the cell and cause mitochondrial damage that, in turn, can lead to decreased ATP production and increased oxidative stress (27, 40). On the other hand, maladaptation in autophagy can result in autophagic cell death (1, 18). In the heart, autophagy is increased in response to aging, starvation and in diseases such as heart failure and ischemia (2, 10, 14, 19). Autophagy activation by mTOR inhibitor has also been reported to prevent hypertrophy (20). However, the role of autophagy in the context of anthracycline cardiotoxicity is a subject of debate (2, 19). Notably, several recent reports have demonstrated that increased autophagy is protective against DOX cardiotoxicity (19, 22). Phosphoinositide 3-kinase γ (PI3Kγ) inhibition has been shown to ameliorate DOX cardiotoxicity by means of enhanced autophagic disposal of DOX-damaged mitochondria (22). Furthermore, DOX has been found to activate a PI3Kγ/Akt/mTOR signaling pathway to promote feedback inhibition of autophagy (22). Interestingly, hearts from mice lacking p38γ, p38δ, or both isoforms have been reported to have low activity of mTOR due to high levels of mTOR inhibitor, DEPTOR (9). Consistent with these studies, our present findings uncovered a decrease in mTOR activity, as evidenced by reduced mTOR-Ser2488 phosphorylation, as well as an increase in LC3-II, a marker of autophagosome formation, in DOX-treated hearts of female p38δ^-/-^ mice that were protected from DOX-induced cardiotoxicity. These results suggest that mechanistically, cardioprotection against DOX seen in female p38δ^-/-^ mice could be linked to enhanced autophagy resulting from a decrease in the mTOR-dependent negative feedback inhibiting the autophagy triggered by DOX.

## LIMITATIONS

This study was designed based on the well-established and widely used acute DOX treatment protocol, and involved an administration of a single dose of DOX (20 or 30 mg/kg) delivered via an i.p. injection to mice, followed by an acute survival monitoring period of 10 days. To better mimic human chemotherapeutic regimens, we intend to employ a chronic DOX treatment regimen in our future studies, in which mice are treated with multiple smaller doses of DOX over a prolonged survival period (22, 50).

Previous studies have described cardiotoxicity elicited by treatment of WT mice with 20 mg/kg DOX in similar acute studies (41, 45). However, we did not observe cardiotoxicity at this dose in 15 week-old mice of any genotype examined, necessitating the usage of a higher dose of DOX (30 mg/kg) in this study. The differential susceptibility to DOX20 could be due to differences in the genetic background and/or age of the mice used in different studies. Our present work showed that, in contrast to 15 week-old mice, 22 week-old female WT mice are in fact susceptible to DOX20-induced cardiotoxicity. Clearly, further investigation is warranted of the impact of age on DOX-triggered cardiomyopathy.

## CONCLUSIONS

In this study, we uncovered the crucial sex-specific role of the p38δ MAPK in anthracycline cardiotoxicity. We report that systemic genetic deletion of p38δ protected female mice against DOX-induced cardiotoxicity. p38δ deletion resulted in improved cardiac mechanical and electrical function and thereby increased rate of survival in DOX-treated female mice. The improved cardiac function correlated with reduced fibrosis and enhanced autophagy.

## ACKNOWLEDGEMENTS

We would like to acknowledge the excellent technical assistance provided by Aaron Koppel, Loubna Al Dammad, and Christian Miccile for this study.

