## Supplementary material for "p38δ genetic ablation protects female mice from anthracycline cardiotoxicity": Online Data Supplement

**Supplemental Table 1. Sex-differences in 5-day survival of mice treated with DOX30**

| Mice | DOX (mg/ml) | Male Survival Rate (%) | Female Survival Rate (%) | Male vs Female p-value |
|---|---|---|---|---|
| Control | 0 | 100 | 100 | 0.99 |
| WT | 30 | 40 | 90 | 0.01* |
| p38$\gamma^{-/-}$ | 30 | 50 | 73 | 0.15 |
| p38$\delta^{-/-}$ | 30 | 50 | 100 | 0.01* |
| P38$\gamma^{-/-}\delta^{-/-}$ | 30 | 40 | 82 | 0.02* |

**Supplemental Figures:**

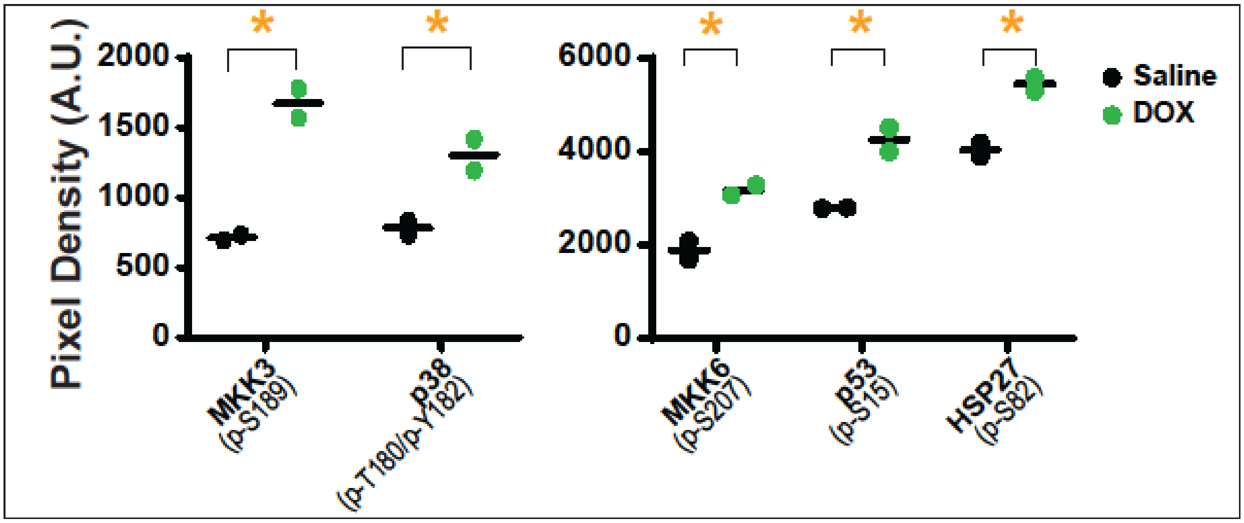


**Supplemental Figure 1. Sustained activation of p38 MAPK pathway by DOX in WT female hearts 5 days post-treatment.** Groups of WT female mice were treated with Saline (n=2) or DOX30 (n=3), and heart tissue lysates were prepared 5 days post-DOX as detailed in the Methods section. Simultaneous detection of the relative levels of the indicated p38 MAPK pathway protein phosphorylation was performed using RayBio C-series Human and Mouse MAPK Pathway Phosphorylation Array Kit (RayBiotech, Inc, Peachtree Corners, GA) according to the manufacturer's instructions. Protein samples from each group were pooled and a total of 600 μg of protein were loaded per each array membrane. Densitometry data were extracted from scanned images of the membranes using ImageJ software in accordance with the manufacturer's guidelines. Briefly, pixel densities of constant sized circle areas of the dots were determined using ImageJ with a plugin after normalizing for the membrane's specific background using the formula recommended by the kit's manual. Since each phosphoprotein was represented by two dots (technical replicates), statistical analysis could be performed on the comparisons. Statistical analysis was performed using two-tailed unpaired t test with GraphPad Prism 6 software. * $p<0.05$

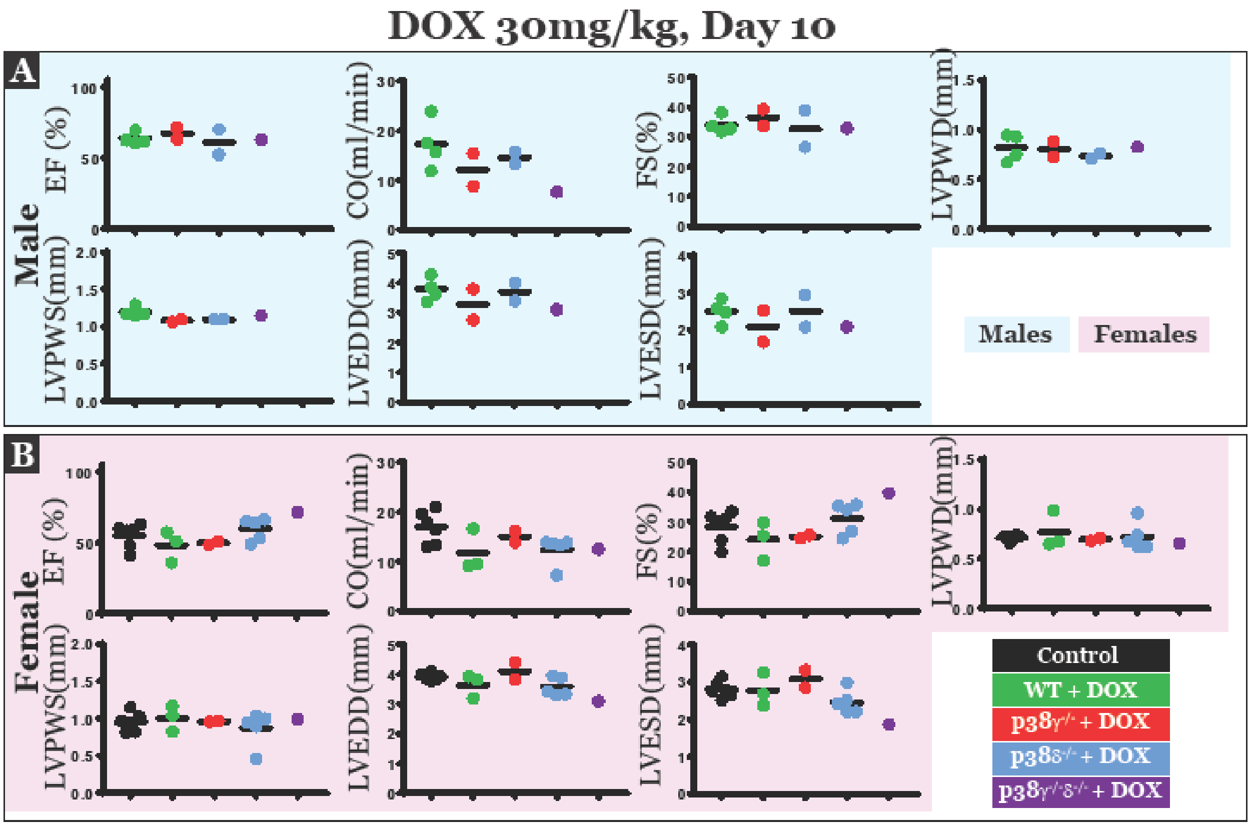


**Supplemental Figure 2. No differences in cardiac mechanical function and LV structure were observed in the surviving mice 10 days post-DOX treatment.** Parameters of cardiac mechanical function such as ejection fraction (EF), cardiac output (CO) and fractional shortening (FS), as well as left ventricular parameters such as left ventricular posterior wall thickness during diastole/systole (LVPWD and LVPWS, respectively) and left ventricular end diastolic/systolic diameter (LVEDD and LVESD, respectively) were measured in male (**A**, upper blue panel) and female (**B**, lower pink panel) mice treated with 30 mg/kg DOX 10 days post-treatment.

**Male, DOX30mg/kg**

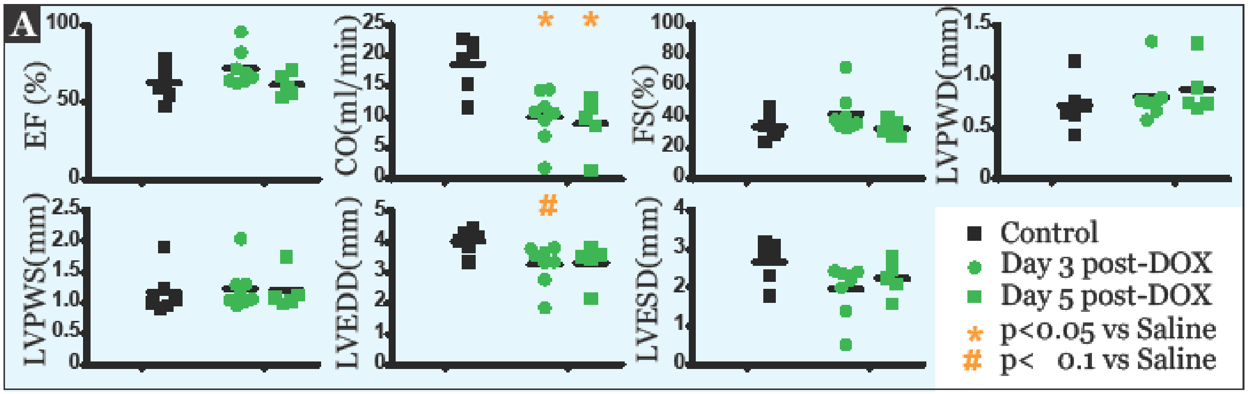


**Supplemental Figure 3. Cardiac function was altered early in the survival period in male WT mice treated with 30 mg/kg DOX.** Echocardiographic parameters were measured on Days 3 and 5 post-DOX treatment in male WT mice. Cardiac output was significantly reduced on Days 3 and 5 with no changes in EF or FS. LV diameter was also reduced on Day 3 post-DOX.

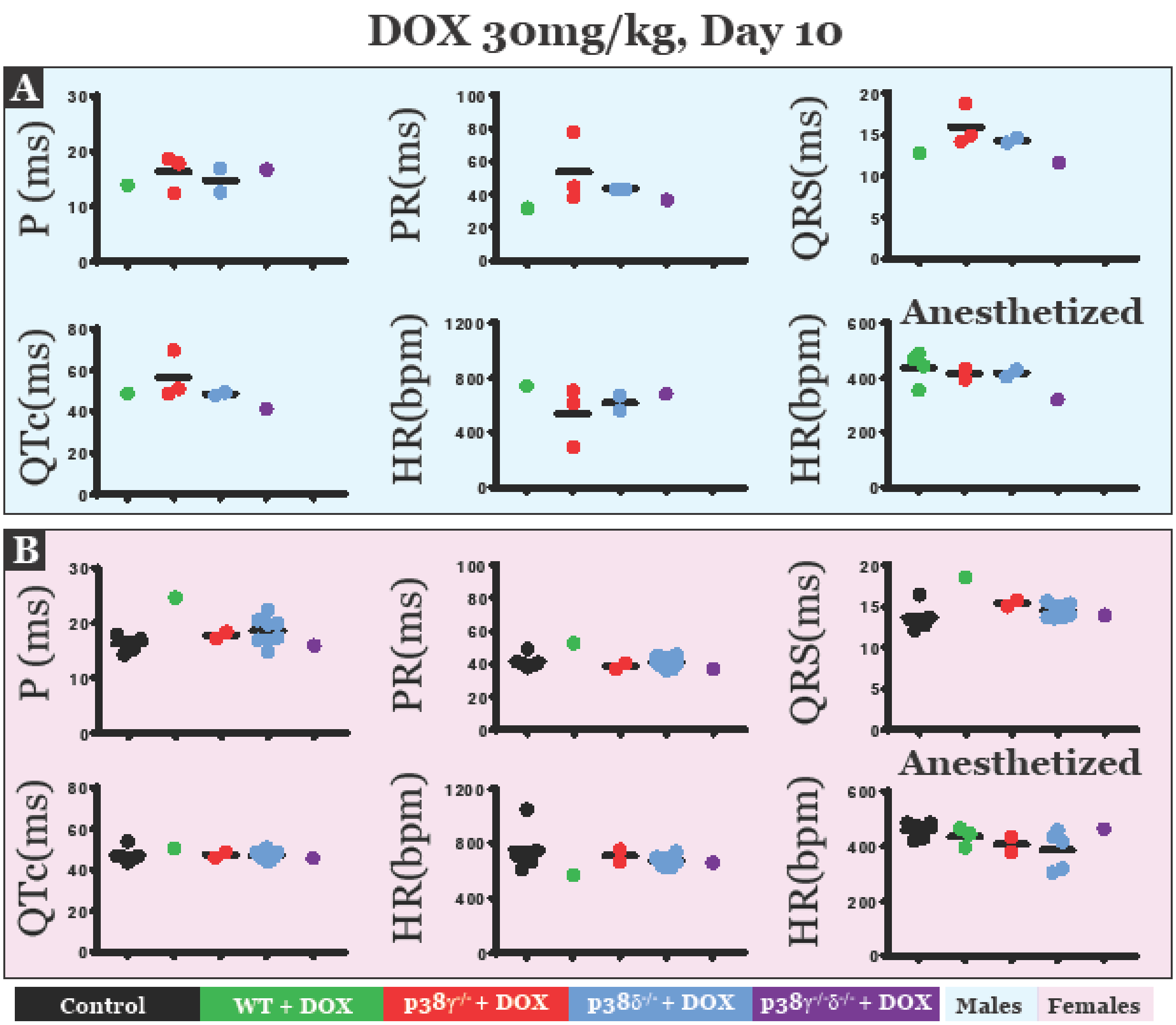


**Supplemental Figure 4. No differences in cardiac electrical function were observed in the surviving mice 10 days post-DOX treatment.** ECG parameters such as the P wave interval (P), P-R interval (PR), QRS duration (QRS), Q-T interval corrected for heart rate (QTc) and heart rate (HR) were measured in conscious male (**A**, upper blue panel) and female (**B**, lower pink panel) mice treated with 30mg/kg DOX 10 days post-treatment. HR in anesthetized mice was also measured for comparison.

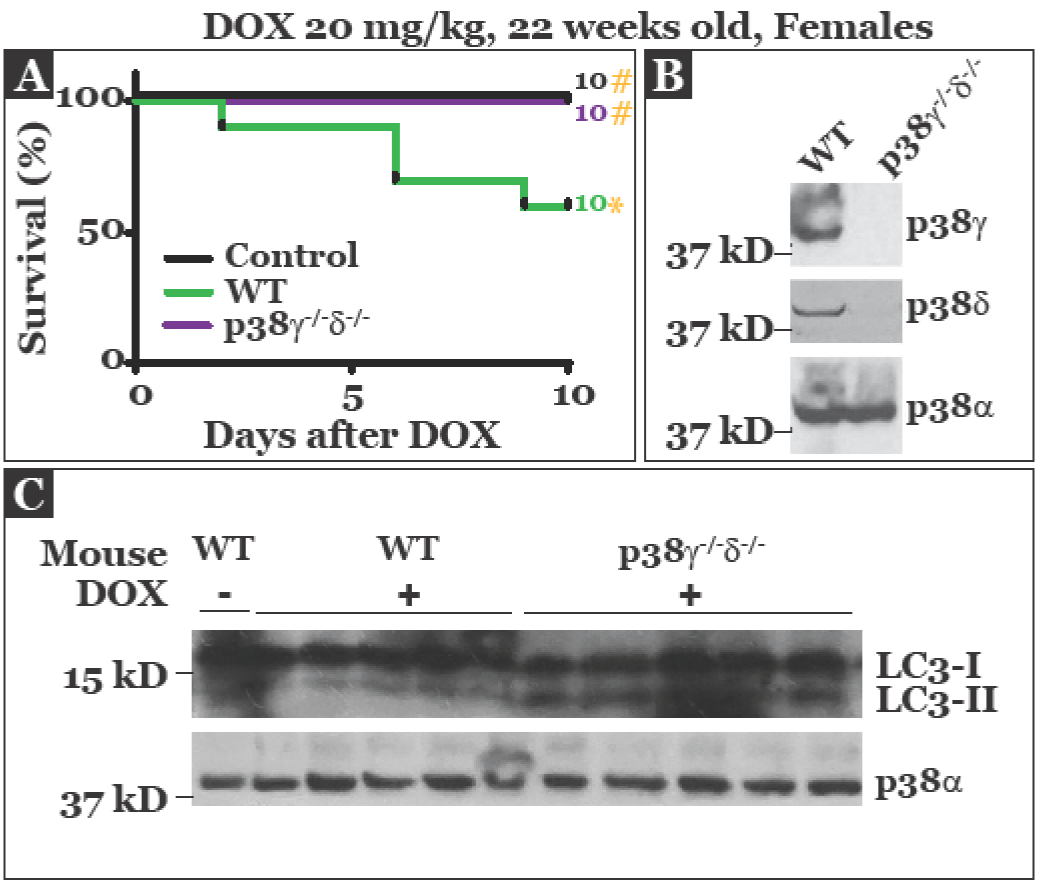


**Supplemental Figure 5. Cardioprotection and improved survival in the 22 week-old female p38γ$^{-/-}$δ$^{-/-}$ mice treated with 20 mg/kg DOX. A)** Kaplan-Meier curves demonstrating a significantly higher survival in 22 week old female p38γ$^{-/-}$δ$^{-/-}$ mice treated with DOX20 relative to female WT mice treated with DOX20 and similar to controls. **B)** Western blots confirming the deletion of p38γ and p38δ proteins in the heart lysates from p38γ$^{-/-}$δ$^{-/-}$ mice. p38α levels were assessed as a loading control. **C)** Increase in expression of the autophagic marker LC3-II was observed in heart tissues from female p38γ$^{-/-}$δ$^{-/-}$ relative to female WT mice treated with DOX20. 50 μg of protein was loaded per lane of the lysates isolated from the hearts of the individual mice treated as indicated in the figure. p38α levels were assessed as a loading control.
